# RIG-I-dependent tumor-intrinsic type I interferon signaling restricts growth in breast cancer 3D culture

**DOI:** 10.1101/2025.08.10.669545

**Authors:** Katherine Liu, Lise Mangiante, Roni Levin-Konigsberg, Cristina Sotomayor-Vivas, Wenting Yang, Kaitlyn Spees, Zhicheng Ma, Jennifer L. Caswell-Jin, Christina Curtis, Michael C. Bassik

## Abstract

Discovery of key growth drivers that can be targeted for therapy is a central goal in cancer research. While high-throughput CRISPR screens have revolutionized our ability to identify gene dependencies in cancer, most large-scale screens are conducted in two-dimensional (2D) culture systems that fail to recapitulate tumor organization and behavior. To uncover architecture-dependent vulnerabilities in breast cancer, we performed parallel CRISPR interference (CRISPRi) screens in 2D and three-dimensional (3D) cultures of MCF7 cells, an estrogen receptor–positive (ER+) breast cancer model representative of a high risk of relapse, luminal subtype. Knockdown of IFNAR2 and TYK2 conferred a growth advantage in 3D cultures, implicating type I interferon signaling as a tumor-intrinsic suppressor of proliferation in 3D spheroids. Transcriptomic and functional analyses demonstrated that type I IFN signaling is endogenously activated in 3D spheroids via RIG-I–mediated sensing of cytosolic double-stranded RNA, leading to TBK1 activation and induction of interferon-stimulated genes (ISGs). This tumor-intrinsic IFN response slowed proliferation in 3D culture, independent of exogenous stimuli or the presence of immune cells. Analysis of bulk, single-cell, and spatial transcriptomic datasets from breast cancer patients revealed that a subset of tumors exhibit elevated IFN signaling in cancer cells, including in immune-depleted tumor cores, consistent with a tumor-intrinsic IFN signature. Our findings uncover an IFN-mediated growth-suppressive program shaped by 3D tumor architecture, and contribute towards a better understanding of the role of tumor-intrinsic IFN activity.

## Introduction

Estrogen receptor-positive (ER+) breast cancer accounts for the majority of breast cancer cases and can recur many years after initial treatment. Integrative genomic analyses have stratified ER+ tumors into multiple molecular subtypes, offering improved prognostic resolution beyond classical clinical and pathological features^1,2,3^. Some subgroups, such as those enriched for copy number gains at 17q23, have been associated with late recurrence and therapy resistance, yet the functional impact of many subtype-defining alterations remains poorly understood.

While CRISPR screens have accelerated the discovery of key cancer vulnerabilities, most large-scale screens are conducted in two-dimensional (2D) monolayer cultures, which lack key aspects of tumor biology such as spatial constraints, mechanical forces, and complex cell-cell interactions^4,5^. Three-dimensional (3D) spheroid models more closely mimic *in vivo* tumor architecture, including cell-cell interaction and localized signaling, all of which can reshape gene regulatory networks and cellular dependencies^6–9^. These insights have particular relevance in breast cancer, where numerous studies have shown that gene expression patterns and drug sensitivities observed in 3D cultures better reflect *in vivo* responses compared to 2D monolayers^10–14^. Supporting this, recent 3D CRISPR screens in lung cancer have revealed drug sensitivities and resistance mechanisms that are not captured in 2D systems, many of which correlate with patient prognosis and have been validated *in vivo*^15^.

Type I IFNs play pivotal roles in tumor immune surveillance, modulation of the tumor microenvironment, and regulation of immune checkpoint pathways. Interferon signaling exhibits a nuanced duality in breast cancers, where activation has been shown to induce apoptosis, but can also contribute to the evolution of drug resistance, immune suppression, and metastasis^16,17,18^. Recent studies suggest that the impact of type I IFN on cancer therapy response is highly variable, and contingent upon the strength, duration, and source of IFN signaling stimuli^19,20^. However, most of the research focus has been on the roles of immune-mediated IFN signaling, whereas the impact of tumor-intrinsic IFN signaling has been less studied.

To identify architecture-dependent regulators of breast cancer growth, we performed parallel CRISPR interference (CRISPRi) screens in 2D and 3D cultures of MCF7, an ER+ breast cancer cell line representative of IC1. This dual-screening approach uncovered a striking 3D-specific growth-promoting phenotype upon knockdown of IFNAR2 and TYK2, core components of the type I interferon (IFN) receptor complex. Transcriptomic profiling of 3D spheroids revealed robust upregulation of interferon-stimulated genes (ISGs), even in the absence of immune cells. We then investigated upstream signaling pathways and identified RIG-I (encoded by *DDX58*), a cytosolic RNA sensor, as the key initiator. RIG-I is best known for detecting viral RNA and triggering innate immune responses, but it is increasingly recognized as a key regulator in cancer, where it modulates tumor-intrinsic stress signaling and acute inflammatory responses^23,24^. In our spheroid model, RIG-I is triggered by endogenous dsRNA, revealing a tumor-intrinsic IFN program shaped by spatial organization and signaling dynamics of 3D tumor architecture. Importantly, this IFN program is not restricted to *in vitro* spheroid models. Analysis of transcriptomic data from breast cancer patients across bulk, single-cell, and spatial datasets consistently showed a subset of tumors with enriched IFN signatures in cancer cells, including within immune-depleted tumor core regions, supporting the presence of a tumor-intrinsic IFN signature *in vivo*.

Together, these findings reveal that type I IFN signaling functions as a growth-constraining program that emerges in 3D tumor models, independent of exogenous stimulation by immune cells. This work expands our understanding of the multifaceted roles type I IFN can play in cancer growth, further highlighting the therapeutic potential of this pathway.

## Results

### CRISPRi screens in 2D vs 3D reveal tumor-intrinsic type I IFN signaling as a critical regulator of growth in 3D breast cancer spheroids

To investigate growth dependencies in ER+ breast cancer, we performed CRISPR-interference (CRISPRi) screens in both 2D monolayers and 3D spheroids of MCF7, a breast cancer cell line representative of high-risk of relapse luminal tumors (Integrative Cluster 1, IC1)(Fig. 1a; Supplementary Fig. 1a-b). MCF7 exhibits genomic features such as amplification of the 17q23 amplicon, which includes a number of putative oncogenic drivers with potential therapeutic relevance. To functionally interrogate these genes and other candidate druggable targets, we designed a custom CRISPRi sub-library targeting candidate drivers from this amplicon, and used it alongside a sub-library that includes gene targets of a large panel of FDA-approved drugs, as well as all known kinases and phosphatases (DTKP library). Strikingly, we found that knockdown of several components of the type I IFN signaling pathway, including IFNAR2 and TYK2, yielded much stronger positive growth phenotypes in 3D compared to 2D cultures (Fig. 1b; Supplementary Fig. 2c). To validate this observation, we performed a competitive growth assay by co-culturing equal numbers of MCF7 cells with control (safe) sgRNA and either sgRNA against IFNAR2 or TYK2 (Fig. 1c-d; Supplementary Fig. 2d). Cells with IFNAR2 or TYK2 knockdown consistently outcompeted control cells in 3D spheroids, significantly more so than in 2D, highlighting a striking growth advantage specifically dependent on 3D tumor architecture.

**Figure 1.**
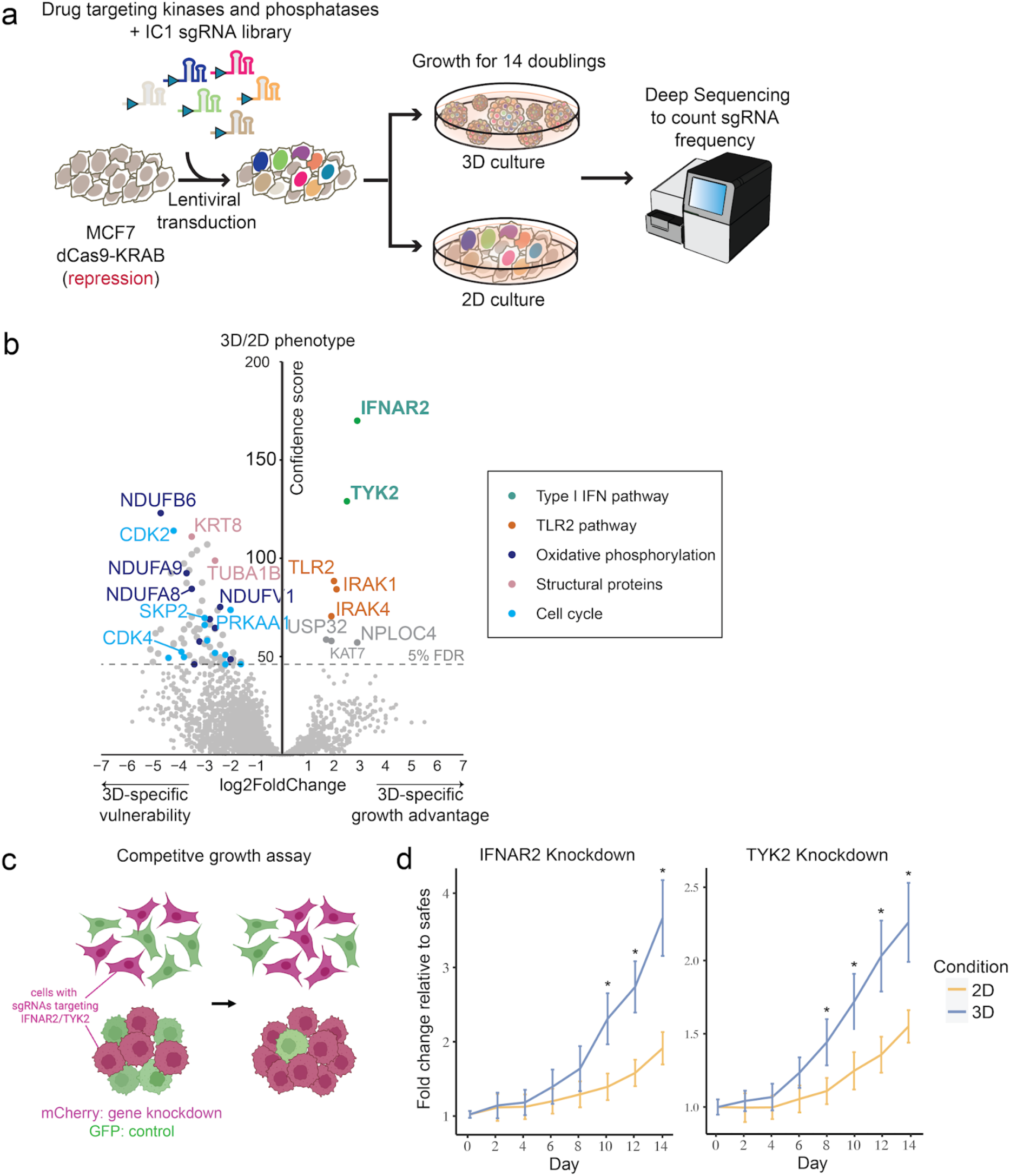
CRISPRi screens in 2D and 3D MCF7 cells identify the type I interferon pathway as a 3D-specific hit. **a)** Schematic of CRISPR-inference (CRISPRi) screen in MCF7 cells aimed at identifying 3D-specific vulnerabilities. **b)** Volcano plot showing genes corresponding to sgRNAs that were significantly enriched in 3D compared to 2D (*n* = 2 independent experiments) using casTLE analysis. Values on the x-axis show the effect size of each gene and the y-axis show confidence score. The dashed line denotes the 5% FDR threshold. Selected hits are colored by pathways: type I IFN signaling (green), TLR2 pathway (orange), oxidative phosphorylation (purple), structural proteins (pink), and cell cycle genes (blue). Full gene-level screen results for 2D and 3D conditions are provided in Supplementary Table 1-3. **c)** Competitive growth assay of MCF7 cells with safe sgRNA and sgRNA against IFNAR2 (left) or TYK2 (right).

Since IFNAR2 and TYK2 are the key receptor and kinase in the type I IFN response, we hypothesized that the type I IFN pathway is upregulated in 3D. We therefore examined the effect of IFNAR2 knockdown on gene expression in cancer cells in 3D and 2D cultures by RNA-seq (Supplementary Fig. 3a). Differential gene expression analysis revealed that genes involved in the type I IFN pathway, such as *OAS*, *ISG15*, and *BST2*, were significantly upregulated in 3D cultures, suggesting activation of the type I IFN pathway in the 3D environment (Fig. 2a; Supplementary Fig. 3b). These results support the hypothesis that the type I IFN pathway is activated in 3D, and normally restricts the growth of MCF7 spheroids. Upon IFNAR2 knockdown, the enrichment of type I IFN response genes was lost (Fig. 2a, right panel; Supplementary Fig. 3c-d), indicating that the induction of the IFN response in 3D is dependent on autocrine or paracrine signaling through the IFN receptor IFNAR2. More detailed analysis of specific stress pathways (Fig. 2b) further highlighted the upregulation of genes involved in JAK-STAT signaling, RNA editing, and nucleic acid sensing, pathways that are all modulated by type I IFN signaling. This was corroborated by GO-term analysis (Supplementary Fig. 3b), which showed significant upregulation of IFN-related immune response in 3D cultures.

**Figure 2.**
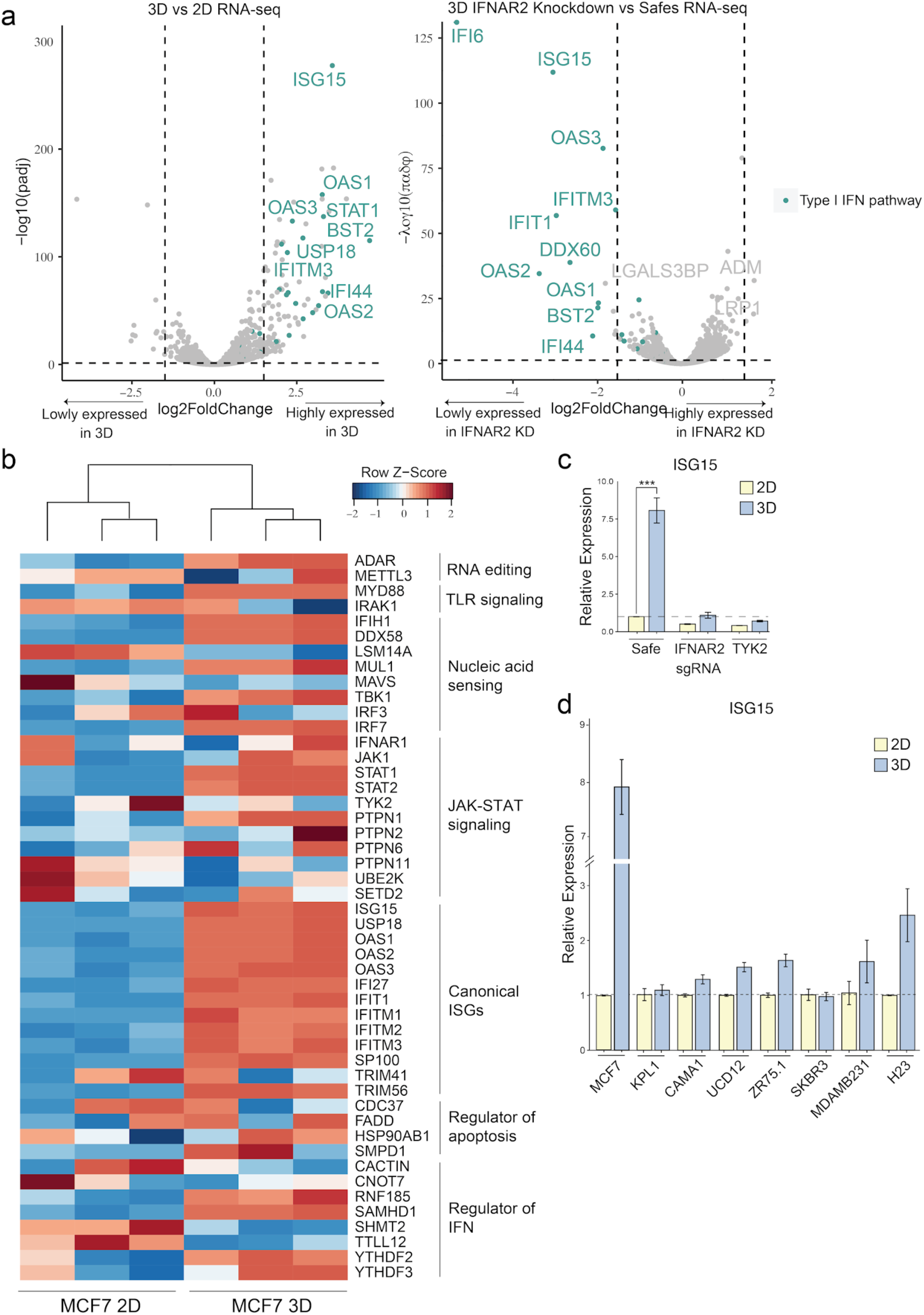
The type I interferon pathway is upregulated in MCF7 3D spheroids. **a)** Volcano plots showing differentially expressed genes in MCF7 3D spheroids compared to 2D culture (left) and MCF7 3D spheroids with IFNAR2 knockdown compared to safe sgRNA control (right). RNA-seq was performed in Illumina NovaSeq 6000, followed by demultiplexing, STAR alignment, making gene-level counts with HTseq, and differential expression analysis with DESeq2. Full differential expression results are provided in Supplementary Table 7. **b)** Heatmap showing the expression of type I interferon ISGs in MCF7 when cultured in 2D or 3D. Cells were cultured in 2D and 3D in triplicates for 4 passages (8 days). Raw z-scores are plotted in the heat map with a focus on type I interferon ISGs based on MSigDB gene set M550. **c)** RT-qPCR results showing *ISG15* expression levels in 2D and 3D MCF7 upon *IFNAR2* or *TYK2* knockdown. **d)** The type I interferon pathway is upregulated in 3D spheroids derived from diverse cell lines.

To further validate these findings, we measured *ISG15* expression levels in MCF7 3D spheroids by RT-qPCR (Fig. 2c). We focused on *ISG15* because it was one of the most strongly upregulated ISGs in our RNA-seq analysis and is a well-established marker of type I IFN signaling^27^. Consistent with the transcriptomic data, *ISG15* expression was significantly elevated in 3D cultures. Moreover, this upregulation was attenuated upon knockdown of IFNAR2 or TYK2, two key components of the type I IFN receptor signaling pathway, supporting that the IFN response in 3D is dependent on canonical receptor-mediated signaling.

To determine the contextual requirements for type I IFN induction in MCF7 cells, we tested whether the upregulation of *ISG15* in 3D cultures could be attributed to secreted factors or the extracellular matrix. Conditioned media collected from 3D spheroids did not induce ISG15 expression in 2D cultures at either 4 or 8 days (Supplementary Fig. 5c), suggesting that soluble factors alone are insufficient to recapitulate the IFN response. Additionally, culturing cells in methylcellulose (MC) media in standard tissue culture plates (2D) did not result in *ISG15* upregulation, whereas spheroids formed in ultra-low attachment plates with MC media exhibited a robust increase in *ISG15* expression (Supplementary Fig. 5a), indicating that 3D context is required beyond matrix exposure alone. We further compared matrix conditions in basement membrane extract (BME) to MC media and observed similarly high *ISG15* levels at day 15 (Supplementary Fig. 5b). This could be because spheroids take longer to form in BME compared to MC.

To assess whether the upregulation of IFN genes is specific to MCF7 cells, we performed similar 2D versus 3D comparisons across multiple cell lines. RT-qPCR results (**Fig. 2d**) revealed elevated *ISG15* levels in 3D spheroids of various cell lines, although the increase was not as pronounced as observed in MCF7 spheroids. Nonetheless, the overall trend remained consistent, reinforcing the hypothesis that 3D culture conditions promote type I IFN signaling across different cell types.

### RIG-I activity induces IFN signaling in 3D spheroids

To further investigate the mechanism of tumor-intrinsic type I IFN production in our 3D cell model, we systematically examined canonical upstream activators of the IFN pathway. The TBK1 (TANK-binding kinase 1) axis represents a central hub for initiating type I IFN responses in an immune cell–independent manner (Fig. 3a). Upon detection of cytosolic nucleic acids, sensors such as RIG-I, MDA5, and cGAS activate TBK1 through MAVS or STING, depending on the type of nucleic acid^28^. Activated TBK1 subsequently phosphorylates and activates the IRF3 and IRF7 transcription factors, which then translocate to the nucleus to drive the expression of type I IFNs.

**Figure 3.**
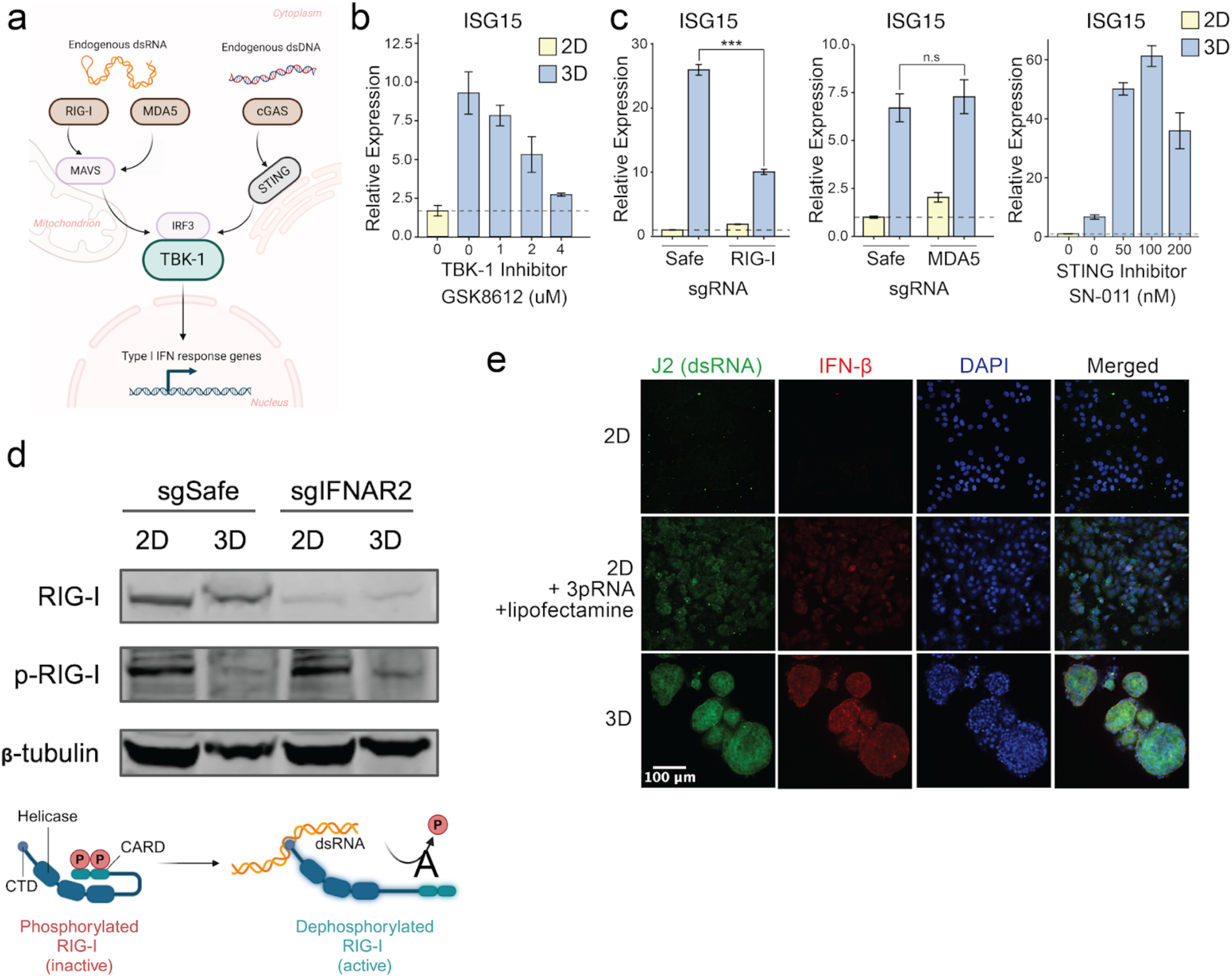
RIG-I activation by dsRNA elicits type I interferon response in MCF7 3D spheroids. **a)** Schematic of cytosolic nucleic acid sensors and the TBK1 pathway. **b)** RT-qPCR results showing ISG15 expression levels in 2D and 3D MCF7 upon TBK-1 inhibition by various concentrations of GSK8612 for 8 days. **c)** RT-qPCR results showing *ISG15* expression levels in 2D and 3D MCF7 upon knockdown or inhibition of cytosolic nucleic acid sensors: RIG-I (left), MDA5 (middle) and STING (right). **d)** Western blot showing phosphorylation state of RIG-I in 2D and 3D MCF7 with sgRNA against safe regions or *IFNAR2*. **e)** Confocal images showing IFN-β and J2 (dsRNA) staining in 2D and 3D MCF7. MCF7 2D cells transfected with triphosphate RNA (3pRNA) using lipofectamine were used as a positive control for J2 staining.

To test whether IFN production in our 3D cell model depends on TBK-1, we treated MCF7 3D spheroids with the TBK-1 inhibitor GSK8612 (Fig. 3b). TBK-1 inhibition led to a concentration-dependent reduction in *ISG15* expression, suggesting that the increased type I IFN production and response may be mediated through the TBK-1 pathway. We then examined whether knocking down upstream sensors of TBK-1, including cytosolic dsDNA sensor STING and dsRNA sensors MDA5 and RIG-I, would impact *ISG15* expression (Fig. 3c; Supplementary Fig. 5d). Among these, only RIG-I knockdown reduced *ISG15* levels, implicating RIG-I as a key upstream activator. Co-treatment with GSK8612 in RIG-I- or IFNAR2-knockdown spheroids did not further reduce ISG15 expression (Supplementary Fig. 5e), consistent with the interpretation that TBK1 acts in concert with RIG-I and IFNAR2 in a shared signaling axis. Unexpectedly, inhibition of STING resulted in an increased IFN response, indicating that STING may not serve as a primary mediator of the 3D-dependent IFN signaling. Further investigation will be required to elucidate the mechanism underlying this unexpected observation.

To better understand how RIG-I is activated in 3D, we next examined its activation status and upstream triggers in spheroid cultures. RIG-I is known to induce type I IFN response upon binding to short cytosolic RNA such as 5’ppp-dsRNAs (3pRNA)^29^. In its inactive state, RIG-I is auto-inhibited by phosphorylation, which inhibits its downstream signaling ability^30^ and upon detection of short cytosolic RNAs, it is dephosphorylated and activated. To assess RIG-I activity in 2D and 3D cultures, we performed immunoblotting. As anticipated, phosphorylated RIG-I was detected in MCF7 2D cells, whereas dephosphorylation of RIG-I was observed in 3D spheroids, indicating RIG-I activation specifically in 3D culture (Fig. 3d). Knockdown of IFNAR2 reduced overall RIG-I protein levels, consistent with the role of IFN signaling in sustaining RIG-I expression through a positive feedback loop. Since double-stranded RNA (dsRNA) is a known activator of RIG-I which leads to type I IFN production, we used confocal microscopy with the J2 antibody, which specifically recognizes and binds to dsRNA structures, to assess dsRNA levels. We observed significantly higher levels of IFN-β and dsRNA in the 3D spheroids compared to 2D cultures (Fig. 3e). This suggests that elevated dsRNA levels in 3D spheroids could be triggering RIG-I activation, leading to type I IFN production.

### Tumor-intrinsic IFN signaling in breast cancer is independent of immune infiltration

Given our observation of IFN signaling in breast cancer 3D spheroids in vitro, we sought to investigate IFN signaling in human patient-derived breast cancer samples. We analyzed bulk and single-cell transcriptomic data from patient tumors. Bulk RNA-seq data from The Cancer Genome Atlas Breast Invasive Carcinoma (TCGA BRCA)^31^ revealed significantly elevated expression of *ISG15*, a marker of type I IFN signaling, in tumor samples compared to normal tissues (Fig. 4a). To assess cell type-specific IFN activity, we next analyzed single-cell RNA sequencing (scRNA-seq) data from a dataset comprised of 119 primary breast tumor biopsy samples from 88 patients across eight publicly available datasets^32^ (Fig. 4b). We calculated an IFN module score using 12 ISGs identified from our RNA-seq of MCF7 3D spheroids (Fig. 2b, Supplementary Table 5) and overlapping with type I IFN gene sets from MSigDB (Supplementary Fig. 6a). In 52 tumors with at least 100 cancer epithelial cells, IFN module scores varied widely across individuals (Fig. 4c). Notably, a subset of tumors exhibited elevated IFN activity, but in general IFN activity did not correlate with immune cell proportion. These data suggest the presence of a tumor-intrinsic IFN signature in some patient tumors that is not always the result of immune cell contact, consistent with our observations in the tumor cell-only 3D spheroid model.

**Figure 4.**
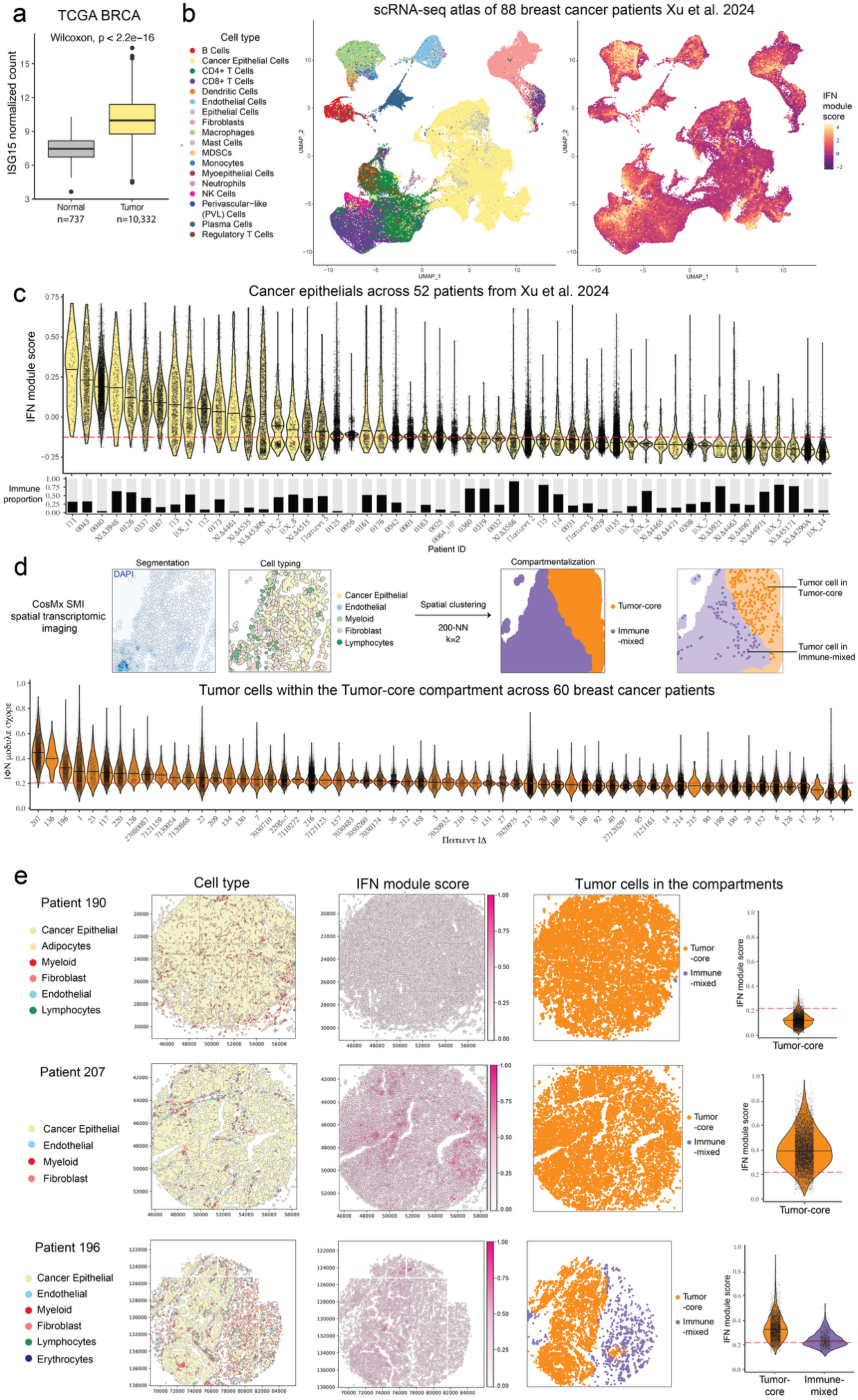
Tumor-intrinsic IFN signaling in breast cancer, independent of immune infiltration. **a)** *ISG15* normalized expression in normal breast tissue (n = 737) versus breast tumor samples (n = 10,332) from the TCGA BRCA cohort. Boxplots show median and interquartile ranges; significance was determined by a two-sided Wilcoxon test. **b)** UMAP visualization of single-cell RNA-seq (scRNA-seq) data of 236,363 cells from 88 breast cancer patients^29^ depicting major cell types (left) and corresponding IFN module scores (right) **c)** Violin plots showing IFN module scores in cancer epithelial cells across 52 patients, each with at least 100 cancer epithelial cells. The red dotted line represents the median IFN module score of all cancer epithelial cells. Bar plots represent the proportion of immune cells for each patient. **d)** Spatial transcriptomics analysis of 450,222 cells from 60 breast cancer patients using CosMx SMI. Representative images show segmentation, cell-type classification, and spatial *k-means* local nearest-neighbourhood (*NN*) clustering based on *200-NN* and *k=2* to identify spatially distinct cell type compartments. Violin plots display IFN module scores in 232,467 tumor cells within the tumor-core compartment across patients. The red dotted line represents the median IFN module score of all tumor cells within the tumor-core compartment. **e)** Spatial maps from three patients illustrating cell-type distributions (left), IFN module scores (middle), and tumor compartmentalization (right), with violin plots comparing IFN module scores in tumor-core and immune-mixed regions. The red dotted line represents the median IFN module score of all tumor cells.

While scRNA-seq offered valuable insight into cell type–specific IFN activity, it lacks spatial resolution and cannot capture the architectural context of gene expression. To overcome this limitation, we analyzed in-house spatial transcriptomic data from 60 breast cancer patients using CosMx SMI, which preserves tissue organization and enables *in situ* mapping of IFN signaling across the tumor landscape. Tumor regions were segmented, classified by cell type, and subjected to spatial *k*-means local nearest-neighbor (NN) clustering based on 200-NN and *k*=2 to define spatially distinct cell type compartments, identifying two distinct transcriptional compartments: tumor-core (orange) and immune-mix (purple) (Fig. 4d, top; Supplementary Fig. 7c). Within these regions, tumor cells in the immune-depleted tumor-core compartment exhibited variable IFN module scores, with a subset of tumors maintaining high IFN activity despite the absence of immune cells, reinforcing the presence of tumor-intrinsic IFN signaling and consistent with our observation of type-I IFN signaling in breast cancer spheroids in vitro. (Fig. 4c, bottom). Spatial maps of representative patients further illustrated these patterns, with tumor-core and immune-mixed compartments showing distinct IFN module distributions (Fig. 4e). For example, the whole tissue section of patient 207 was classified as a tumor-core compartment. Despite the dense packing of tumor cells and limited immune infiltration, high levels of IFN signaling were observed across the whole sample.

Collectively, these findings indicate that IFN signaling in breast cancer can be tumor-intrinsic and does not always correlate with immune infiltration. The variability observed across patients underscores the need to further investigate the regulatory mechanisms driving tumor-intrinsic IFN activation and its implications for tumor progression and immune responses.

## Discussion

Here we investigated architectural regulators of cancer growth using parallel CRISPRi screens in MCF7 2D monolayers and 3D spheroids, and identified core components of the type I IFN receptor complex as growth suppressors in 3D. Transcriptomic analyses revealed robust induction of ISGs under 3D conditions, a response that was consistently observed across multiple cell lines. Mechanistic investigation demonstrated that this IFN response is initiated by RIG-I, a cytosolic RNA sensor, which detects dsRNAs enriched in 3D cultures and activates downstream TBK1 signaling. This tumor-intrinsic IFN signature is also evident in patient tumors, where subsets of cancer cells, particularly within immune-depleted tumor cores, exhibit elevated ISG expression. These results highlight type I IFN signaling as a tumor-intrinsic pathway that is selectively engaged in 3D tumor models, independent of external stimuli.

Using microscopy, we detected the accumulation of cytosolic dsRNA specifically in MCF7 3D spheroids, but not in 2D cultures (Fig. 3e). Several mechanisms may contribute to this selective enrichment of dsRNA. One possibility is the reactivation of endogenous retroelements (ERVs), which are known to generate immunostimulatory dsRNA intermediates when de-repressed^33,34^. Recent studies have shown that circular RNAs (circRNAs), especially when misprocessed or forming double-stranded structures, can activate RIG-I in cancer cells^35^. Given their abundance and dysregulation in tumors, circRNAs may contribute to the accumulation of immunostimulatory RNA species in 3D spheroids, triggering tumor-intrinsic IFN signaling. A recent study by^36^ has shown that oncogenic transformation in breast cancer can suppress innate immune sensing pathways, including those mediated by RIG-I, implying that cytosolic RNA sensing represents a barrier that tumor cells must overcome or adapt to during progression. Thus, the accumulation of dsRNA in 3D cultures may result from increased transcriptional or metabolic stress, reactivation of repetitive elements, or defects in RNA processing. These are all conditions that can generate endogenous ligands capable of activating RIG-I in the absence of viral infection. Future work will be required to elucidate the origins of the tumor cell-intrinsic IFN signaling across diverse cancer types.

While we observed IFN signaling in immune-depleted tumor cores in a subset of patient samples, it was not uniformly elevated in all cases, highlighting interpatient variability (Fig. 4c, 4e). This heterogeneity underscores the need for further investigation into the molecular determinants that govern tumor-intrinsic IFN activation and how it influences tumor growth and immune interactions, as well as the functional consequences of activating this pathway *in vivo*. This observation also raises important questions about how and why this pathway persists in certain tumors despite its antiproliferative effects. In ER+ breast cancer, Shimada et al. found that high ER-signaling (ERS) is associated with reduced type I IFN pathway activity and HLA expression, leading to diminished antigen presentation and impaired T cell infiltration within the tumor microenvironment^37^. Recent work from our group found that ER+ high-risk tumors, which often exhibit lower ER expression levels, harbor extrachromosomal DNA (ecDNA) and show increased IFN signaling^3^. These findings suggest that ecDNA may contribute to the observed IFN elevation in some ER+ tumors. While RIG-I is classically recognized as a cytosolic RNA sensor, emerging evidence suggests that ecDNA accumulation can trigger type I IFN responses through the production of potential source of functional RNAs, such as microRNAs, small interfering-like RNAs, and short hairpin-like RNAs^38,39^, potentially linking ecDNA to innate IFN activation observed in these patients and our 3D models. Further investigation is needed to determine whether ecDNA is enriched in IFN-high ER+ breast tumors, and to assess whether ecDNAs engage RIG-I sensing pathways, thereby contributing to tumor-intrinsic type I IFN activation.

Although 17q23-amplified genes did not rank among the top depleted hits in either 2D or 3D screens, we observed a subtle trend toward depletion (Supplementary Fig. 2a-b). These observations raise the possibility that amplification of this locus contributes to a cumulative fitness advantage through the action of multiple genes, rather than conferring dependency on a single dominant driver. Further work will be required to identify how multi-gene networks may contribute to growth of amplicon-driven tumors.

Given the growth-suppressive effects of type I IFN signaling, we expected that tumors would downregulate or evade this pathway during progression. Intriguingly, we detected persistent tumor-intrinsic IFN signaling in established tumors, suggesting that those tumors may instead acquire mechanisms to tolerate or adapt to persistent IFN activity. Our findings align with earlier studies reporting that elevated type I IFN levels can paradoxically correlate with both tumor progression and therapeutic resistance^19,39–41^. The complexity of the role of IFN in cancer progression is further highlighted by the study of^18^, which described how these cytokines can either promote anti-tumor immunity or facilitate immune evasion, depending on the context. In tumors where IFN signaling remains transient and controlled, it enhances immune priming and supports T cell infiltration. However, when IFN signaling is sustained, it can shift the tumor environment towards immunosuppression, reinforcing the tumor’s resistance to immune therapies. Recent studies from Reticker-Flynn et al. have shown that persistent tumor-intrinsic type I IFN signaling can promote immune tolerance and accelerate lymph node metastasis, highlighting a pro-tumor role for IFN signaling in cancer progression^42^. This dual role of IFN may help explain why elevated IFN levels are observed in certain tumors, where prolonged IFN exposure may drive an immunosuppressive state rather than sustaining anti-tumor activity.

Our findings underscore the importance of studying tumor-intrinsic immune pathways within physiologically relevant 3D models. By uncovering a type I IFN response that constrains tumor growth *in vitro* yet persists in subsets of immune-depleted patient tumors, our study raises fundamental questions about how cancer cells modulate or tolerate stress-associated signaling programs. Whether this persistence reflects incomplete pathway suppression, a reprogrammed IFN state with altered downstream consequences, or an adaptive strategy that enables survival under selective pressure remains to be determined. Even so, given certain limitations of ER+ breast cancer mouse models—such as species-specific differences in estrogen signaling, challenges in modelling long-latency distant relapse, and difficulty to reflect the physiology of postmenopausal women in which most ER+ tumors arise^43,44^—complementary *in vitro* 3D systems offer a tractable platform for mechanistic investigation. Moving forward, dissecting the temporal dynamics, epigenetic regulation, and downstream effectors of tumor-intrinsic IFN signaling may offer critical insights into the balance between tumor suppression and immune evasion. Such knowledge could inform more nuanced therapeutic strategies that consider not only the presence of IFN activity but its cellular source, duration, and functional context. The ability to model key aspects of this response *in vitro* provides a unique opportunity to dissect the induction and regulation of the tumor-intrinsic IFN response in the presence and absence of infiltrating immune cells.

## Methods and Materials

### Cell lines

The MCF7-dCas9-KRAB cell line was a generous gift from the Howard Chang lab (Stanford University). Cells were maintained in Dulbecco’s Modified Eagle Medium (DMEM; Gibco #11995073) supplemented with 5% fetal bovine serum (FBS; Gibco #A5670701), 1% penicillin-streptomycin (PS; Gibco #15-140-63), and 10 µg/mL blasticidin (Gibco #A1113903) to maintain selection for dCas9-KRAB expression. Cells were cultured at 37 °C with 5% CO₂ in a humidified incubator and regularly tested to ensure they were mycoplasma-free.

### 3D spheroid cultures

For the preparation of 1L of 0.75% methylcellulose media, 7.5 g of methylcellulose (Fisher Scientific #M352-500) and a magnetic stir bar were placed in a sterile glass bottle, then autoclaved using a dry cycle. After cooling to room temperature, the following components were added: 5% FBS (Gibco #A5670701), 1% PS (Gibco #15-140-63), 1% L-glutamine (Gibco #25030081), and DMEM (Gibco #11995073) to a final volume of 1L. The mixture was stirred at 4°C overnight or until the methylcellulose was fully dissolved. The prepared medium was stored at 4°C for short-term use (up to 1 month).

### CRISPR-dCas9 Interference Screening in 2D and 3D MCF7 Cells

For CRISPR-dCas9 interference screening, we utilized the Drug Targeting Phosphatases and Kinases sgRNA library to repress target genes in MCF7 cells cultured as 2D monolayer and 3D spheroids. Briefly, 10 sgRNAs were designed for each gene, with safe-targeting sgRNAs serving as negative controls. Cells were transduced with the library at 0.3 MOI and grown for 14 passages. To assess growth phenotypes, the frequencies of each sgRNA in the cell population before and after growth were compared. sgRNA abundance was determined via deep sequencing to identify genes that influence cell growth under the given conditions.

### Calculation of growth phenotype

To measure the growth phenotype, casTLE was used to quantify the relative fitness of cells carrying different sgRNAs^43^. After 14 passages, genomic DNA was harvested from the cell population, and sgRNA sequences were amplified by PCR. The amplified products were then subjected to deep sequencing on the NextSeq 550 platform to determine the relative abundance of each sgRNA. Raw sequencing data were processed and aligned to the sgRNA library to calculate the frequency of each sgRNA in the population before and after growth. sgRNA frequencies from both 2D and 3D conditions were also analyzed to identify 3D-specific hits.

### Competitive growth assay

MCF7 cells expressing the indicated sgRNAs were seeded into either standard tissue-culture treated (for 2D monolayer culture) or ultra-low attachment (for 3D spheroid culture) 24-well plates. The competitive growth assay was adapted from Han et al., 2020, with modifications to support 3D spheroid culture and longitudinal analysis by flow cytometry. Approximately 50,000 mCherry-labeled cells expressing gene-targeting sgRNAs were mixed with 50,000 GFP-labeled cells expressing a safe sgRNA, and co-seeded into wells, in triplicates. Cultures were maintained for 14 days and passaged every 2 days, with spheroids gently dissociated using Accutase before passaging.

At each time point, cells were dissociated into single-cell suspensions and analyzed on an Attune NxT Flow Cytometer (Thermo Fisher Scientific #A24858) to quantify the proportions of GFP- and mCherry-positive cells. The mCherry:GFP ratio at each time point was normalized to the ratio at day 0 to calculate fold changes over time. To account for baseline effects of marker expression and competition, fold changes were further normalized to a control mixture of mCherry- and GFP-labeled cells both expressing the safe sgRNA. Relative growth phenotypes for each sgRNA were calculated separately for 2D and 3D conditions, and 3D/2D growth ratios were computed to assess the context specificity of each gene perturbation.

### RNA-seq experiments and analysis

MCF7 cells expressing control (safe) or *IFNAR2* sgRNA were cultured as 2D monolayer or 3D spheroids. RNA was extracted with TRIzol (Invitrogen #15596018) and purified with RNeasy Mini kit (Qiagen #74104). Quality control and RNA sequencing was performed by Novogene using Illumina NovaSeq 6000 using a 150 bp paired-end sequencing protocol. Sequencing reads were aligned to the GRCh38 genome reference using STAR 2.7^46^. Gene-level counts were generated using HTseq 2.0^47^. Differentially expressed genes of the conditions were analyzed using DESeq2 1.44^48^.

### RT-qPCR

MCF7 cells expressing control (safe) or *IFNAR2* sgRNA were cultured as 2D monolayer or 3D spheroids. RNA was extracted with TRIzol (Thermo Scientific #15596018) and purified with RNeasy Mini kit (Qiagen #74106). cDNA synthesis was performed using SuperScript™ IV Reverse Transcriptase (Thermo Scientific #18090050). qPCR was performed using the Applied Biosystems™ TaqMan™ Fast Advanced Master Mix for qPCR (Thermo Scientific #4444557) on CFX Connect Real-Time PCR Detection System (BioRaD), with appropriate TaqMan™ primers (Supplementary Table 8).

### Confocal immunofluorescence imaging

For 2D cell cultures grown on coverslips, cells were fixed in 4% paraformaldehyde in PBS for 15 min at room temperature. Cells were washed twice with PBS and then permeabilized with 0.2% Triton X-100 in PBS for 15 minutes at 4 °C, followed by blocking in 1% FBS in PBS for 30 minutes at room temperature. Coverslips with cells were then incubated with the primary antibodies at a dilution of 1:100 for 1 hour at room temperature. After three PBS washes, the coverslips with cells were incubated with the secondary antibodies at a dilution of 1:2,500 and 1 µg/mL DAPI for 1 hour at room temperature, followed by three PBS washes. Imaging was done using a spinning-disk confocal microscope (Eclipse Ti, Nikon, CSU-W1, Yokogawa) with a 20x objective.

### Immunoblotting

Cells were lysed in RIPA buffer supplemented with phosphatase and protease inhibitor cocktails (Thermo Scientific #78440). The lysates were incubated on ice for 30 minutes and then clarified by centrifugation at 16,000g for 10 minutes at 4 °C. Protein concentration was measured using the Bradford assay, and lysates were prepared with NuPAGE™ LDS Sample Buffer (4X) (Invitrogen #NP0007). Membranes were probed with antibodies against RIG-I (Cell Signaling #3743), phosphorylated RIG-I Thr170 (Thermo Fisher Scientific #BS-5300R) and Beta-tubulin (Cell Signaling #86298), at a dilution of 1:1,000. Secondary antibodies, anti-rabbit and anti-mouse IRDye-conjugated antibodies from Fisher Scientific, were used at a dilution of 1:10,000 (no. NC9401841, NC9401842, NC0110517, and NC9030091). Membranes were imaged using an Li-Cor Odyssey CLx. Uncropped western blot images can be found in Supplementary Fig. 6b.

### TCGA BRCA analysis

*ISG15* expression levels were assessed in normal and tumor samples from the TCGA BRCA dataset. Raw RNA-seq expression data for ISG15 were downloaded from the TCGA Data Portal. Data were processed and normalized using the TCGAbiolinks R package. Differential expression of *ISG15* between normal and tumor samples was evaluated using the limma package, with log2-transformed normalized counts. Statistical significance was determined using an adjusted p-value threshold of <0.05. Visualizations, including boxplots and heatmaps, were generated using the ggplot2 R package.

### Single-Cell RNA-Seq analysis

To assess tumor-intrinsic IFN signaling in human breast cancer, we analyzed publicly available single-cell RNA-sequencing (scRNA-seq) data from Xu et al., 2024, a breast cancer atlas comprising 119 tumor biopsies from 88 patients. Processed data were downloaded and integrated into a Seurat v4 object for downstream analysis. Cells were filtered based on canonical marker expression and annotations provided by the authors. We focused specifically on cancer epithelial cells, identified using epithelial markers (e.g., EPCAM) and curated cell type labels provided by the authors. To quantify type I IFN signaling activity, we used the AddModuleScore function in Seurat to calculate IFN module scores for each cell. The module consisted of interferon-stimulated genes (ISGs) commonly associated with type I IFN signaling (Supplementary Table 5). These genes were selected based on the top DEGs from Figure 2B and MSigDB-defined interferon response gene sets. Module scores were computed across all cancer epithelial cells, and patients with fewer than 100 epithelial cells were excluded from patient-level analyses to reduce sampling bias. Inter-patient variability in IFN module scores was visualized using violin plots, and module scores were compared with immune cell proportions to assess the extent of immune-independent IFN signaling. All data processing and visualization were performed using R (v4.2.1) and Seurat (v4.3.0)^49^ .

### Spatial transcriptomic profiling using CosMX platform

Spatial transcriptomic data were generated internally using the NanoString CosMX Spatial Molecular Imager (SMI) platform on formalin-fixed paraffin-embedded (FFPE) tissue sections. Tissue processing, probe hybridization (CosMX Human 6000 Gene Panel), subcellular imaging, and initial transcript quantification were performed by colleagues in our laboratory according to the manufacturer’s standard protocol (NanoString Technologies). Briefly, the CosMX platform employs multiplexed in situ hybridization and high-resolution iterative fluorescence imaging to quantify transcript abundance at subcellular resolution. Image registration, segmentation, and transcript localization were performed using NanoString’s image analysis pipeline, and processed data were provided as cell-by-gene expression matrices with associated spatial coordinates and metadata. All downstream analyses were performed using custom R scripts and Seurat (v4.3.0). Cells with fewer than 50 transcripts or had overlaps with other cell nuclei were excluded from analysis. Gene expression matrices were normalized by total transcript counts per cell using Seurat’s SCTransform method V2. Integration of 8 sequencing batches were integrated using Seurat’s anchor-based approach. Cell type annotation was guided by canonical marker gene expression using GeneVector (v0.0.1)^50^. Following the methods described in Schürch and colleagues^51^, a 200-nearest neighbor (200-NN) graph in physical (x–y) space using Euclidean distances between cells was used to build a neighborhood composition matrix. Spatial compartmentalization was then assessed using k-means clustering with *k* = 2, applied independently within each cell’s local 200-NN graph, to identify spatially distinct subclusters. Spatial projection plots were generated using the physical x–y coordinates provided by the CosMX platform for each segmented cell. Using Python SpatialData (v0.4.0)^52^, we visualized cells in their native spatial segmentation, colored by cell type annotation, or expression of specific genes or gene signatures. Plots were generated using custom scripts in Python (v3.10) with libraries including scanpy, matplotlib, and seaborn. For spatial compartment assignment visualization, k-means cluster labels derived from the 200-NN neighborhood graph (with k = 2) were overlaid onto the tissue coordinate map, allowing identification of spatially distinct compartments within each cell type. Spatial plots were rendered at high resolution to preserve single-cell localization, and representative tissue regions were selected for inclusion in figures based on cell density, signal quality, and biological relevance.

## Data Availability

Raw and processed sequencing data generated in this study, including CRISPRi screen counts and RNA-seq expression matrices, will be deposited in the Gene Expression Omnibus (GEO) under accession number [to be provided upon acceptance]. CosMx imaging data will be made available through AtoMx (Bruker), a cloud-based, fully-integrated informatics platform for spatial biology that enables advanced analytics, global data sharing and collaboration. Previously published datasets used for reference or comparison are cited in the Methods section. All other relevant data are available from the corresponding authors upon reasonable request.

## Code Availability

Key code dependencies and parameters are described in the Methods. Custom scripts used for CRISPRi screen analysis (casTLE processing, volcano plots), RNA-seq differential expression, and spatial transcriptomics clustering are available from the corresponding authors upon request.

## Acknowledgements

This work was supported by the National Cancer Institute U54 grant (U54CA261719) to C.C and M.C.B.. We thank Joanna Watson for insightful discussions and feedback throughout the course of this study. We also thank members of the Curtis Lab, including Katie Houlahan, Marni McClure and Clemens Weiss. We are grateful to Grace Chen for her valuable insights on RIG-I and generous help with mass spectrometry. We also thank Howard Chang for generously providing the MCF7 CRISPRi cell line.

## Contributions

Conceptualization: K.L., M.C.B and C.C.

Cell culture: K.L., W.Y.

CRISPR library design: K.L.

CRISPR library cloning: K.S.

Screens and experiments: K.L.

RNA-seq data generation and analysis: K.L.

Confocal imaging: K.L., R.L.-K.

Single-cell RNA-seq analysis: K.L., L.M., C.S.-V.

Survival analysis: K.L., K.E.H

Spatial transcriptomic patient cohorts: J.C-J., C.C.

Spatial transcriptomic data generation: Z.M.

Spatial transcriptomic analysis: K.L., L.M.

Statistical analyses: K.L.

Supervision: M.C.B. and C.C.

Writing (original draft): K.L., M.C.B. and C.C.

Writing (review and editing): all authors.

## Supplementary Figures

**Supplemental Figure 1.**
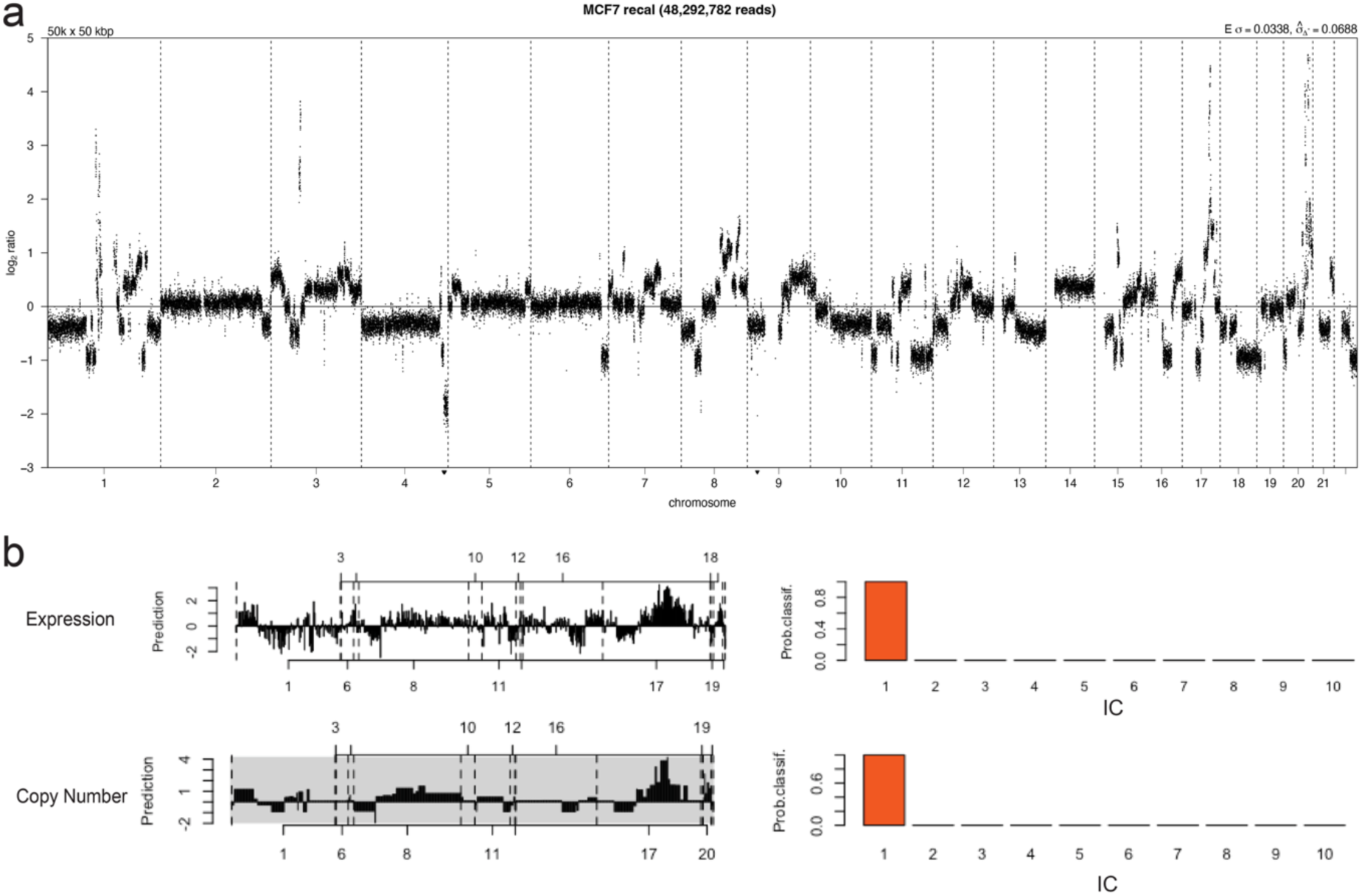
Copy number landscape and integrative subtype classification of MCF7. **a)** Genome-wide copy number profile of MCF7 derived from low-pass whole-genome sequencing (WGS), plotted as log₂ read depth ratios across chromosomes. **b)** Integrative Cluster (IntClust/iC10) classification of MCF7 using the *iC10* R package based on gene expression data (top) and copy number data (bottom). Both modalities assign MCF7 to Integrative Cluster 1 (IC1) with high classification probability.

**Supplementary Figure 2.**
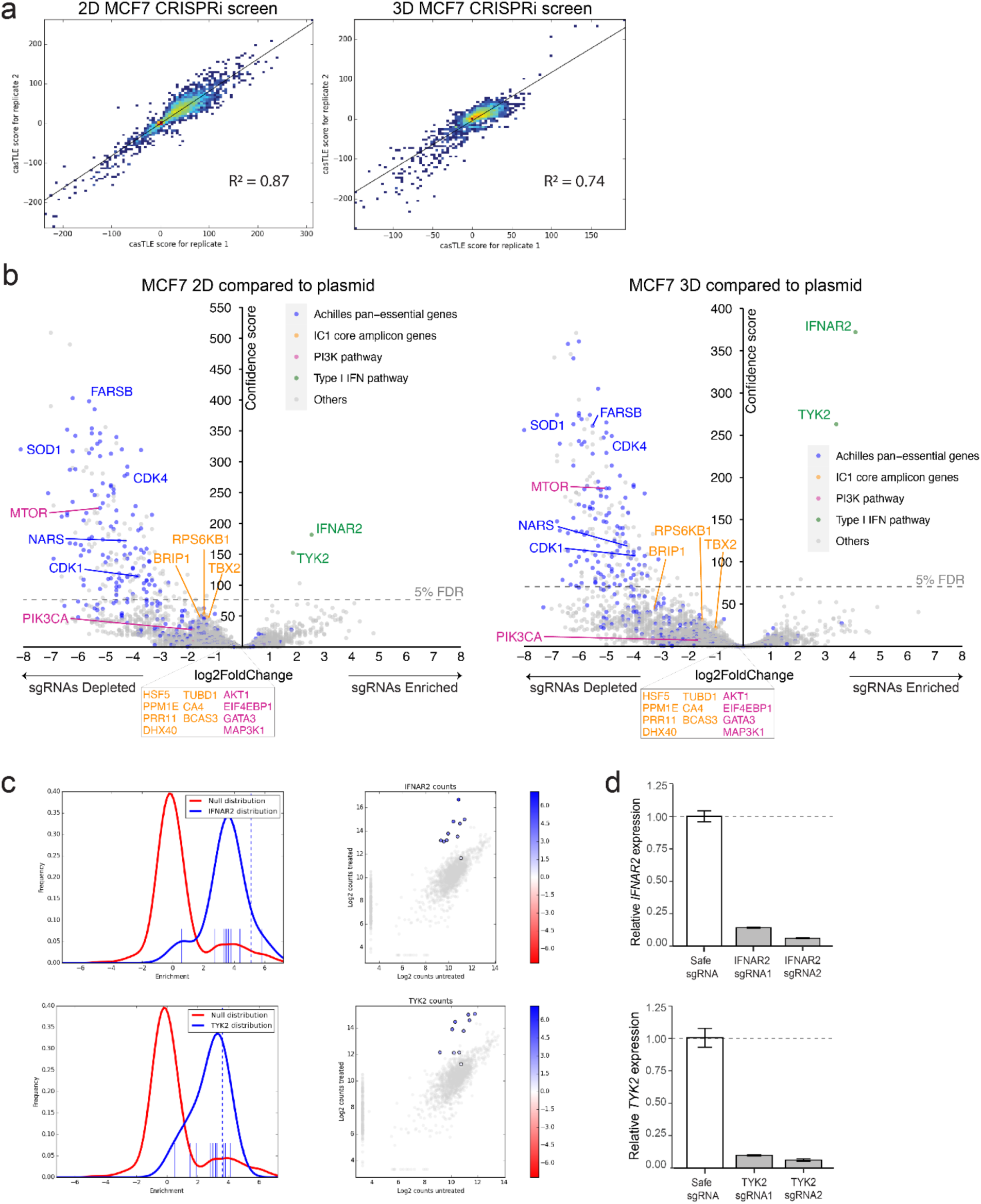
CRISPRi reproducibility, primary screen results, and validation of IFNAR2 and TYK2 knockdown. **a)** Reproducibility of CRISPRi screen replicates in MCF7 cells. Scatterplots show the correlation of casTLE scores between biological replicates for 2D (left) and 3D (right) conditions. Pearson correlation coefficients (R²) are shown for each screen. **b)** Volcano plots of CRISPRi screen results in MCF7 cells cultured in 2D and 3D. Each plot shows genes corresponding to sgRNAs significantly enriched or depleted in Passage 9 compared to plasmid DNA (n = 2 independent experiments), analyzed using casTLE. The x-axis represents effect size, and the y-axis shows the confidence score. The dashed line denotes the 5% FDR threshold. Blue points represent Achilles pan-essential genes, orange points indicate genes within the IC1 core amplicon, pink points indicate PI3K pathway genes, and grey points denote all others. *MTOR*, a PI3K pathway gene, is also classified as pan-essential. Selected well-known growth regulators, IC1 genes, PI3K pathway genes, and type I IFN pathway genes are labeled. Full gene-level screen results for 2D and 3D conditions are provided in Supplementary Table 1-2. **c)** Statistical enrichment and read count distributions for IFNAR2 (top) and TYK2 (bottom) sgRNAs. Left: casTLE enrichment score distributions for each gene (blue) compared to the null distribution (red). Right: Scatterplots of log₂-transformed sgRNA counts in untreated versus treated samples, highlighting sgRNAs targeting IFNAR2 and TYK2 (colored by casTLE effect size). **d)** RT-qPCR validation of knockdown efficiency for *IFNAR2* (top) and *TYK2* (bottom) in MCF7 cells transduced with two independent sgRNAs per gene. Expression is normalized to safe sgRNA control. Bars represent mean ± SD from technical replicates.

**Supplementary Figure 3.**
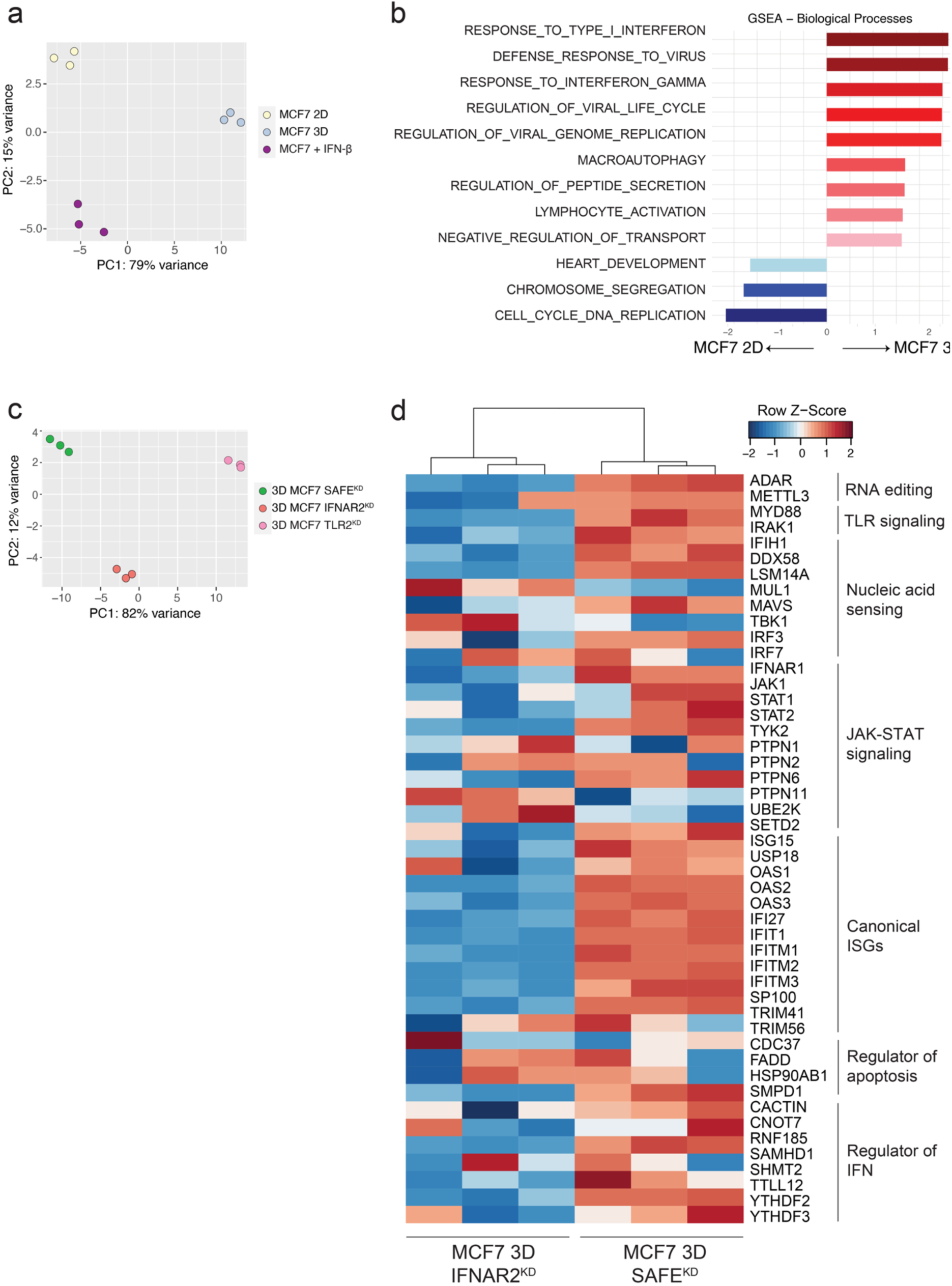
Transcriptomic profiling of MCF7 cells reveals activation of type I IFN signaling in 3D spheroids and its dependence on IFNAR2. **a**) Principal component analysis (PCA) of RNA-seq data from MCF7 cells cultured in 2D, 3D, or treated with recombinant IFN-β (10 ng/mL) for 30 minutes in 2D. Each point represents an individual replicate. **b**) Gene Set Enrichment Analysis (GSEA) of biological processes comparing MCF7 cells grown in 3D versus 2D culture. Selected top-ranked gene sets are shown along with their normalized enrichment scores. **c**) PCA of RNA-seq data from MCF7 3D spheroids with safe control, IFNAR2 knockdown, or TLR2 knockdown. **d**) Heatmap of RNA-seq expression (row Z-scores) for a selected panel of interferon-related genes described in Fig. 2B, shown in MCF7 3D spheroids with IFNAR2 or SAFE sgRNAs.

**Supplementary Figure 4.**
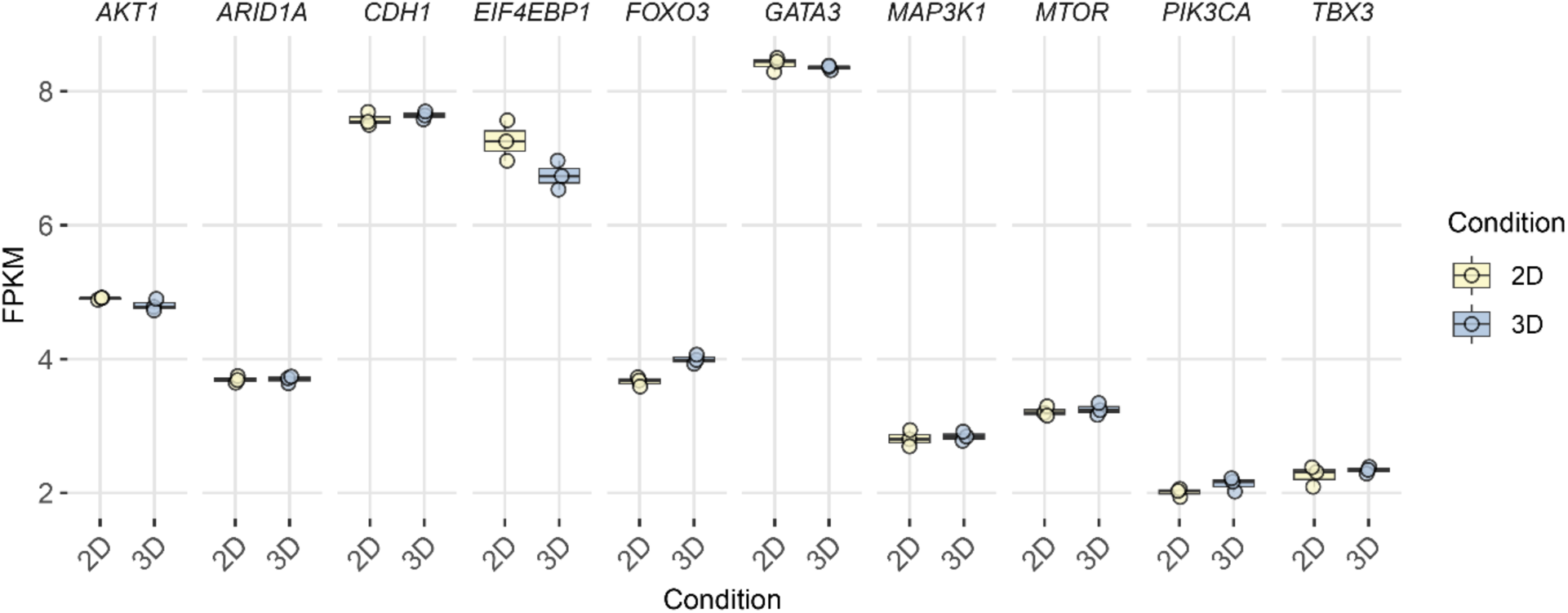
Expression of selected PI3K pathway genes in 2D vs 3D MCF7 cultures. Shown are log-transformed FPKM values for key genes in the PI3K and mTOR signaling pathway in MCF7 cells cultured in 2D or 3D. Each dot represents an independent replicate. Box plots indicate the median and interquartile range.

**Supplementary Figure 5.**
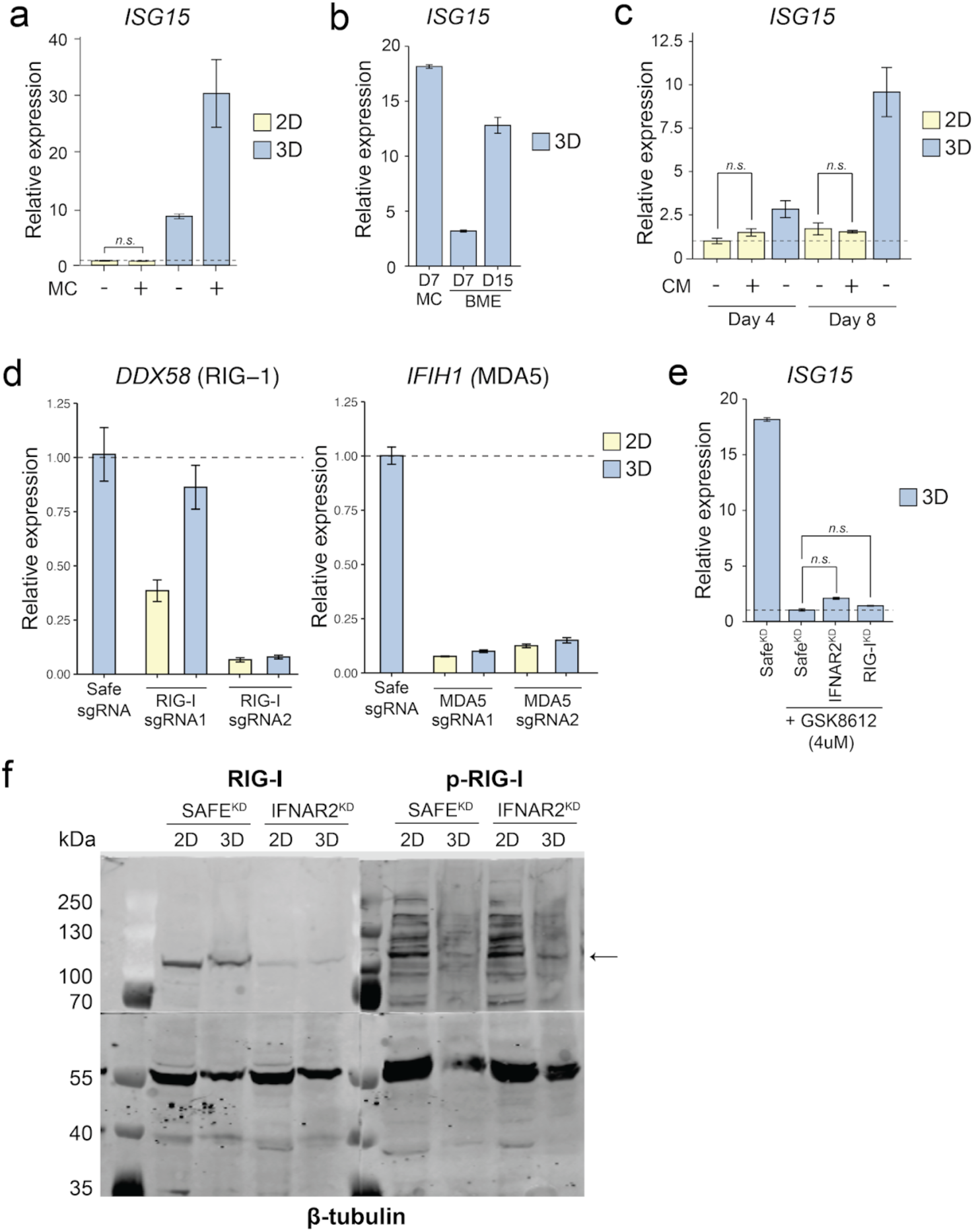
*ISG15* expression and RIG-I pathway activation in MCF7 cells under different culture and perturbation conditions. **a)** RT-qPCR measurement of *ISG15* expression in MCF7 cells cultured in 2D or 3D with or without methylcellulose (MC). The 2D cultures were plated on standard tissue culture plates, while 3D spheroids were grown in ultra-low attachment plates. **b)** *ISG15* expression in MCF7 spheroids cultured in methylcellulose (MC) or basement membrane extract (BME) plates at Day 7 and Day 15. **c)** *ISG15* expression in MCF7 cells cultured in 2D or 3D for 4 or 8 days, with or without conditioned media (CM) collected from 3D spheroids. CM was added to 2D cultures on Day 0 and refreshed with each media change. **d)** *DDX58* (RIG-I) and *IFIH1* (MDA5) knockdown validation compared to safe sgRNA. The more efficient sgRNAs, RIG-I sgRNA2 and MDA5 sgRNA1, were used in Figure 3C. **e)** *ISG15* expression in 3D MCF7 spheroids following treatment with the TBK1 inhibitor GSK8612 (4 μM), with or without knockdown of IFNAR2 or RIG-I. sgRNAs targeting IFNAR2 or RIG-I were introduced via CRISPRi, and gene expression was measured after 8 days in culture. **f)** Uncropped western blot analysis of total RIG-I, phosphorylated RIG-I (p-RIG-I), and β-tubulin (loading control) in MCF7 cells with sgRNA for SAFE or IFNAR2 cultured in 2D or 3D. Arrow denotes p-RIG-I band. Error bars represent mean ± SD from technical replicates. “n.s.”: not significant (unpaired two-tailed t-test).

**Supplementary Figure 6.**
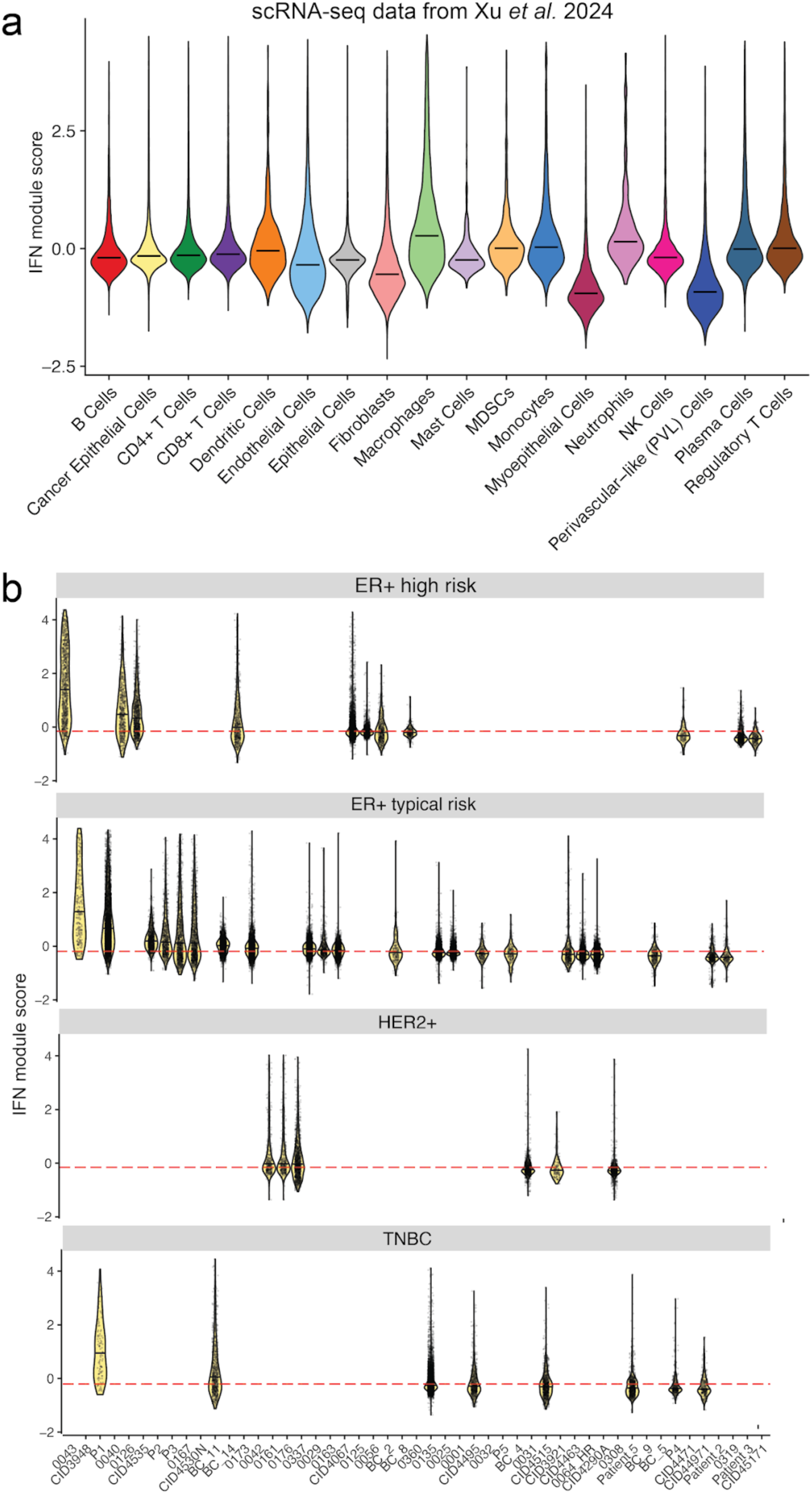
IFN module scores across cell types and breast cancer subtypes in scRNA-seq data from Xu et al. 2024. **a)** Violin plots showing type I interferon (IFN) module scores across annotated cell types from the Xu et al. scRNA-seq dataset. **b)** Violin plots of IFN module scores in cancer epithelial cells, stratified by molecular subtype and recurrence risk category. Each plot represents individual patient tumors within the indicated subtype (ER+ high risk of relapse, ER+ typical risk or relapse, HER2+, and triple-negative breast cancer [TNBC]). Red dashed lines indicate median IFN module scores across all cancer epithelial cells.

**Supplementary Figure 7.**
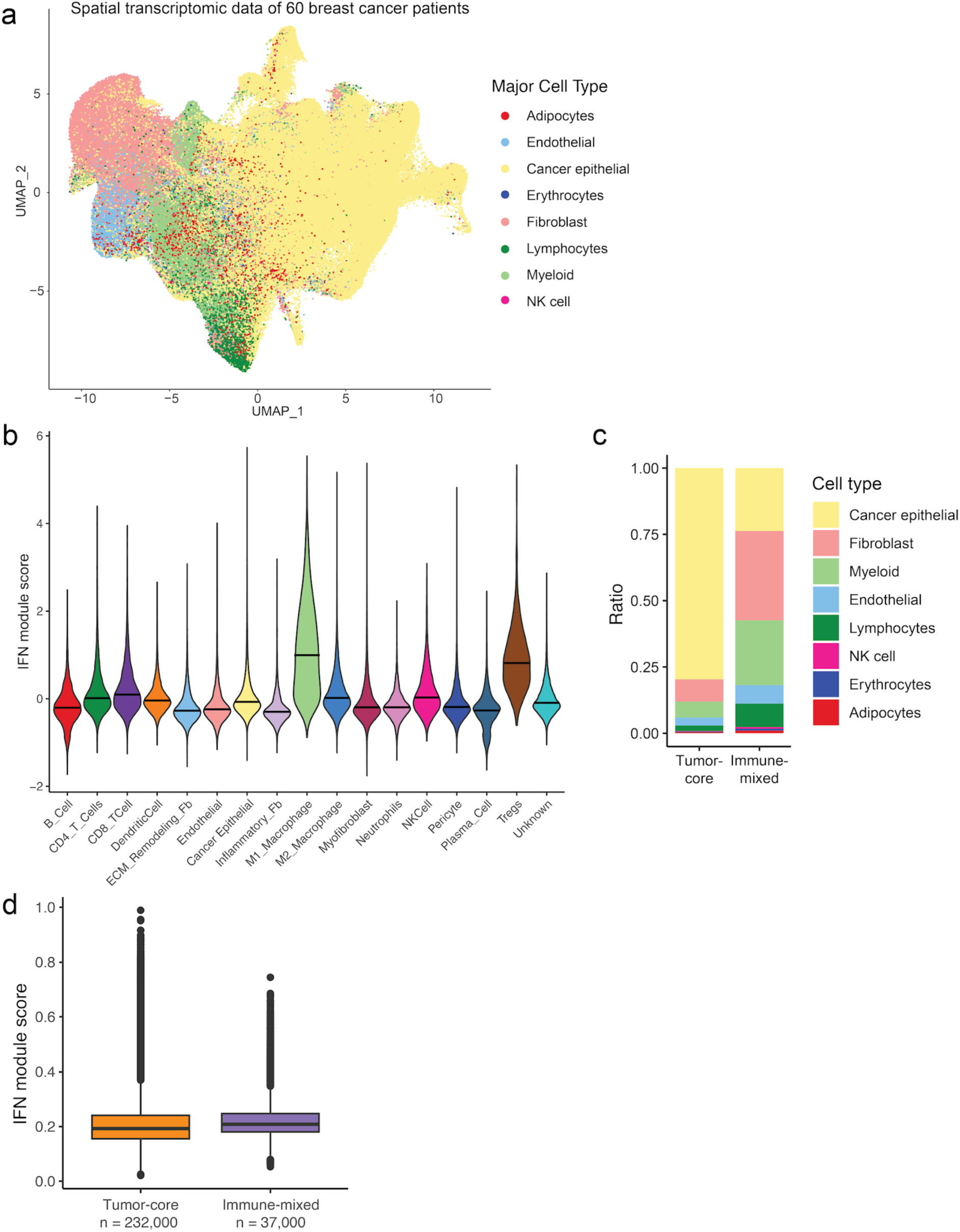
Spatial transcriptomic analysis of IFN module activity in 60 breast cancer patients. **a)** UMAP embedding of 450,222 cells from spatial transcriptomic profiling, colored by major cell type. **b)** IFN module scores across individual annotated cell types, revealing heterogeneity in IFN signaling activity. **c)** Proportion of major cell types in tumor-core (291,854 cells) versus immune-mixed (158,368 cells) spatial compartments. **d)** Violin plots of IFN module scores in cancer epithelial cells across patient samples, stratified by spatial compartment (tumor-core in orange, immune-mixed in purple), with 232,467 and 33,868 cells respectively. Red dashed lines indicate median IFN module scores across all cancer epithelial cells.

